# A neuro-intestinal circuit controls mitochondrial dynamics and stress resistance

**DOI:** 10.1101/2023.06.19.545634

**Authors:** Rebecca Cornell, Wei Cao, Bernie Harradine, Ava Handley, Roger Pocock

## Abstract

Neurons coordinate inter-tissue protein homeostasis to systemically manage cytotoxic stress. Specific neuropeptidergic signals coordinate the systemic mitochondrial stress response (UPR^mt^), but whether chemical neurotransmitters also influence this process is unclear. Here, we show that gamma-aminobutyric acid (GABA) inhibits, and acetylcholine (ACh) promotes the UPR^mt^ in the *Caenorhabditis elegans* intestine. GABA controls the UPR^mt^ by regulating extra-synaptic ACh release through metabotropic GABAB receptors GBB-1/2. We find that elevated ACh levels in animals that are GABA-deficient or lack ACh-degradative enzymes induce the UPR^mt^ through ACR-11, an intestinal nicotinic α7 receptor. This neuro-intestinal circuit is critical for nonautonomously regulating mitochondrial dynamics and organismal survival of oxidative stress. These findings establish chemical neurotransmission as a crucial regulatory layer for nervous system control of systemic protein homeostasis and stress responses.

**One-Sentence Summary:** Regulation of mitochondrial health with neurotransmission.

## Main Text

Maintaining systemic mitochondrial function is crucial for organismal health and survival. Mitochondrial stress responses are essential molecular mechanisms that maintain metabolic homeostasis. Local stress responses can be communicated to distal tissues to systemically combat challenges, and thereby increase the chance of organismal survival. The nervous system is critical for coordinating stress responses across multiple tissues, yet mechanisms by which this is achieved are incompletely understood (*1-7*). In this study, we aimed to better understand how the nervous system maintains systemic mitochondrial function and health.

Neurotransmission is a core communication mechanism of the nervous system. To identify neurotransmitters that mediate systemic mitochondrial health, we screened neurotransmitter synthesis and transport mutants for changes to intestinal expression of the *hsp-6p::gfp* reporter, a well-established readout for the mitochondrial unfolded protein response (UPR^mt^) (*8-12*). We screened loss-of-function mutations affecting dopamine (*cat-2*), octopamine and tyramine (*tdc-1*), serotonin (*tph-1*), glutamate (*eat-4*), gamma-aminobutyric acid (GABA) (*unc-25*) and acetylcholine (ACh) (*cha-1*) signalling (Figure 1A-B) (*13-18*). We found that loss of UNC-25/GAD, a glutamic acid decarboxylase required for GABA synthesis (Figure 1C), induced intestinal *hsp-6p::gfp* expression (Figure 1A-B). Resupplying *unc-25* cDNA to GABA expressing neurons rescued intestinal *hsp-6p::gfp* expression levels in *unc-25(-*) animals (Figure 1D), confirming a role for UNC-25 in nonautonomous regulation of the intestinal UPR^mt^. Animals lacking UNC-25 also showed increased intestinal expression of another mitochondrial stress marker; *hsp-60p::gfp* (Figure S1A-B). Similarly, *sod-3p::gfp* expression – a DAF-16/FOXO transcription factor regulated gene with roles in combatting oxidative stress in mitochondria – was also induced in *unc-25(-*) animals (Figure S1C-D), consistent with previously published data (*19, 20*). We found that UNC-25 loss did not affect *hsp-16*.*2p::gfp*, a reporter for cytosolic heat stress, suggesting that GABA signalling specifically targets mitochondrial stress responses (Figure S1E-F). UPR^mt^ induction is regulated by ATFS-1, a mitochondrial unfolded protein response transcription factor, and the histone modulators UBL-5 and DVE-1 (*10, 21*). Using RNA-mediated interference (RNAi), we silenced *atfs-1, ubl-5* and *dve-1* in *unc-25(-*) animals and found that intestinal *hsp-6p::gfp* induction was suppressed (Figure S1G-H). Therefore, GABA signalling requires canonical transcriptional regulators to activate the intestinal UPR^mt^.

**Fig. 1.**
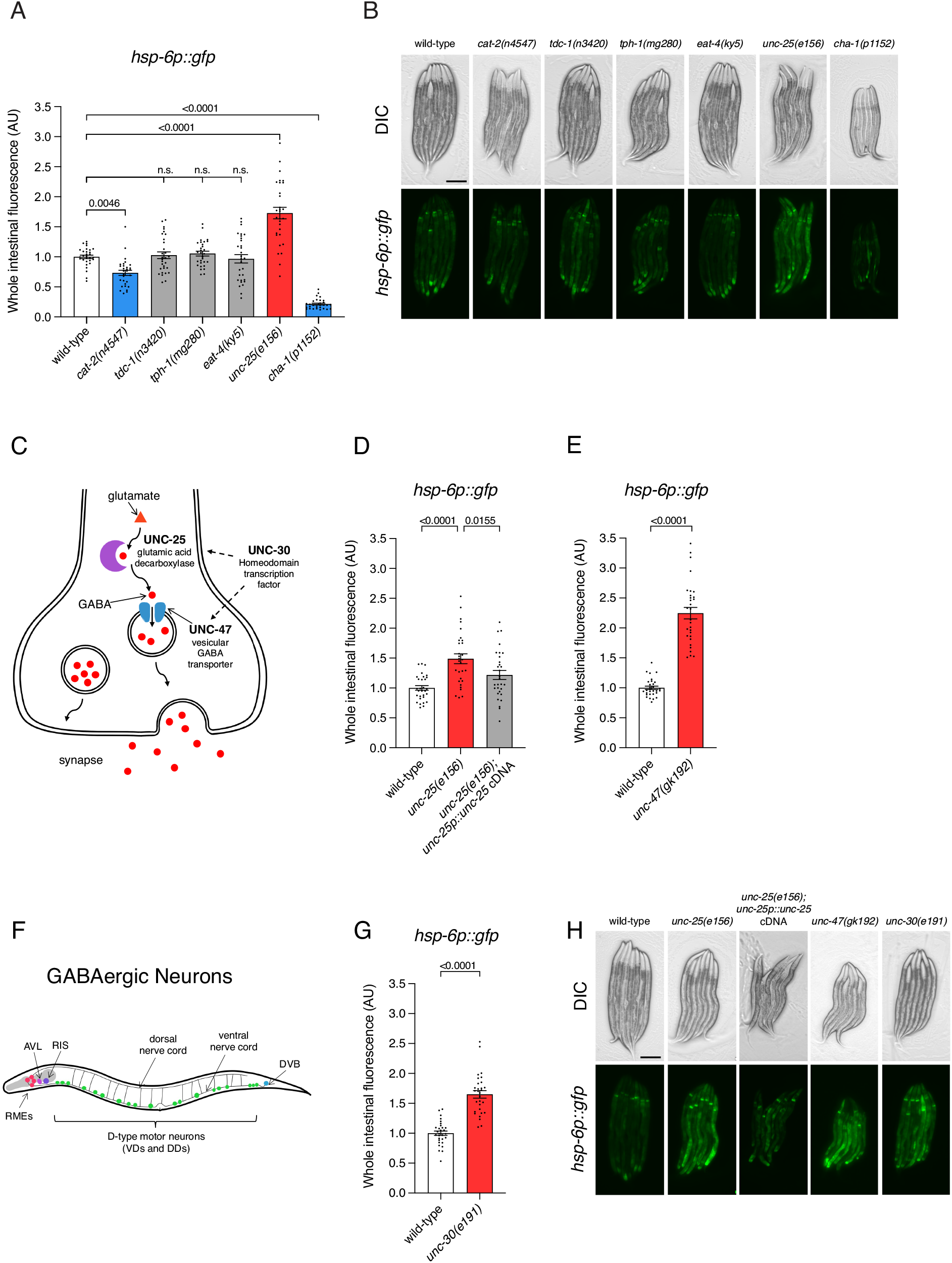
Gamma-aminobutyric acid (GABA) inhibits the cell nonautonomous mitochondrial unfolded protein response (UPR^mt^). (**A** and **B**) Quantification (A) and DIC/fluorescent micrographs (B) of UPR^mt^ reporter (*hsp-6p::GFP*) expression in L4 larvae of wild-type and mutant animals affecting neurotransmitter synthesis and transport. (**C**) Schematic of GABA synthesis (UNC-25/GAD), transport (UNC-47/VGAT), release, and transcriptional regulation (UNC-30/Pitx2). Schematic of GABA synthesis (UNC-25/GAD), transport (UNC-47/VGAT), release, and transcriptional regulation (UNC-30/Pitx2). (**D, E, H**) Quantification (D to E) and DIC/fluorescent micrographs (H) of UPR^mt^ reporter (*hsp-6p::GFP*) expression in L4 larvae of wild-type, *unc-25(e156*) and *unc-25(e156); rpEx2310[unc-25p::unc-25 cDNA]* animals, *unc-47(gk192*), and *unc-30(e191*) animals. (**F**) Schematic of GABA expressing neurons in *C. elegans*. (**G** and **H**) Quantification (F) and DIC/fluorescent micrographs (H) of UPR^mt^ reporter (*hsp-6p::GFP*) expression in L4 larvae of wild-type and *unc-30(e191*) animals. n = 30. *P* values assessed by one-way analysis of variance (ANOVA) Tukey’s post hoc test (A and D) and unpaired t test with Welch’s correction (E and F). Error bars indicate SEM. n.s., not significant. Scale bars, 250μm.

To confirm the role of GABA signalling in controlling the intestinal UPR^mt^, we analysed other components of the GABA signalling pathway (Figure 1C, E-H). The vesicular GABA transporter UNC-47/VGAT is required for packaging GABA into pre-synaptic vesicles for synaptic release (*22, 23*). We found that loss of *unc-47* induces both *hsp-6p::gfp* and *hsp-60p::gfp* expression (Figure 1E,H and S1I-J). The UNC-30/Pitx2 transcription factor is a terminal selector of D-type (DD/VD) GABAergic ventral nerve cord motor neuron identity, where it directly promotes *unc-25* and *unc-47* expression (Figure 1C and F) (*24, 25*). We found that *unc-30* loss also induced *hsp-6p::gfp* expression (Figure 1G-H), suggesting that GABA signalling from the D-type motor neurons represses the UPR^mt^ in the *C. elegans* intestine.

We next sought to determine how GABA influences mitochondria in distal tissues. Several GABA related processes regulate life and health-span, which are concepts closely related to mitochondrial health (*26, 27*). In *C. elegans*, GABA signalling transmits longevity signals through DAF-16/FOXO (*19*). An intermediate component in this pathway is the PLCβ homologue, EGL-8 (*28*). We found that loss of *egl-8* did not induce intestinal *hsp-6p::gfp* expression (Figure S2A,C), suggesting that GABA signalling regulates the intestinal UPR^mt^ and longevity through different mechanisms. We wondered if GABA was acting as a metabolite to influence mitochondrial health through the GABA shunt, where GABA is degraded to succinic semialdehyde (SSA) by GABA-transaminase (GTA-1) and then joins the TCA cycle (*29*). However, loss of the *C. elegans* GABA transaminase, GTA-1, did not influence intestinal *hsp-6p::gfp* expression (Figure S2B-C).

We speculated that GABA signalling mediates the UPR^mt^ from the D-type motor neurons by signalling to downstream GABA receptors. We therefore screened available GABA receptor mutants for *hsp-6p::gfp* induction, focusing on those receptors previously implicated in life and health-span (*20*). We found that loss of the inhibitory UNC-49 and excitatory EXP-1 ionotropic GABAA receptors did not influence the intestinal UPR^mt^ (Figure 2A-C) (*30, 31*). However, loss of both components of the metabotropic GABAB receptor – GBB-1 and GBB-2 – induced intestinal *hsp-6p::gfp* expression (Figure 2A-C). Intestinal *hsp-6p::gfp* expression in the *gbb-2(-); gbb-1(-*) compound mutant was not significantly different to either single mutant, indicating that both components of the GABAB receptor are required in the same pathway to control the UPR^mt^ (Figure 2A-C). Likewise, intestinal *hsp-6p::gfp* expression in the *unc-25(-); gbb-1(-*) compound mutant was not additive compared to either single mutant, showing that GABA synthesis and the metabotropic GABA receptors act within the same genetic pathway to control the systemic UPR^mt^ (Figure 2D). Canonically, GBB-1 and GBB-2 act in concert to reduce neuronal excitability, however, previous studies found that GBB-1 can act independently to influence longevity through DAF-16/FOXO (*19*). This is supported by our data, where *sod-3p::gfp* expression – a readout for DAF-16 activity – is regulated by GBB-1, and not GBB-2 (Figure S3). These data support a role for GABA in UPR^mt^ activation independent to its role in longevity. Single cell sequencing data shows that *gbb-1* and *gbb-2* are expressed primarily in neurons (*32*). Therefore, we resupplied *gbb-1* cDNA under the pan-neuronal *rgef-1* promoter in *gbb-1(-*) animals and found that this rescued the increased intestinal *hsp-6p::gfp* levels (Figure 2E), confirming that the GABAB receptor complex acts in neurons to influence intestinal UPR^mt^ activation.

**Fig. 2.**
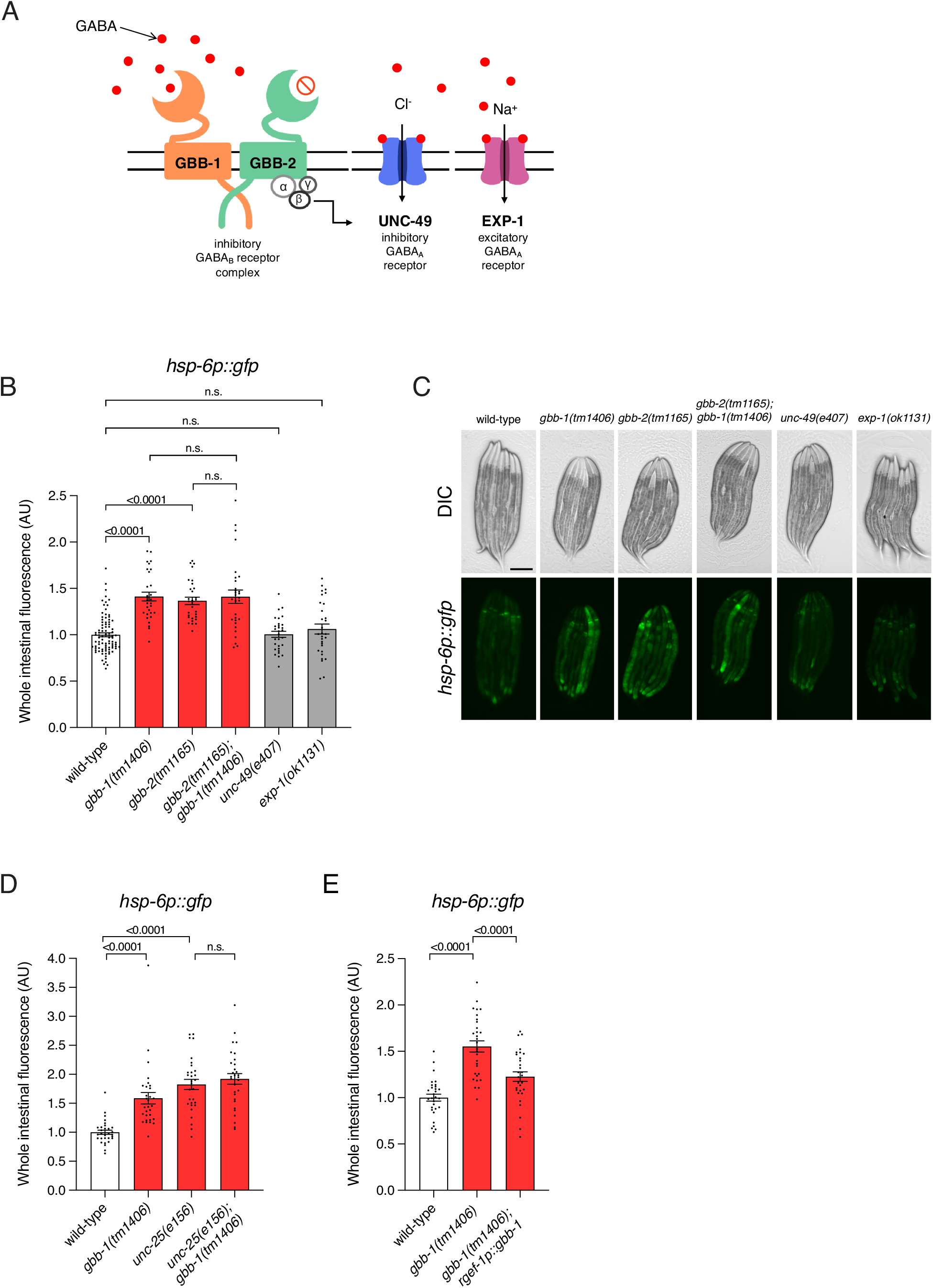
Metabotropic GABAB receptors coordinate cell nonautonomous UPR^mt^. (**A**) Schematic of ionotropic GABAA (UNC-49 (inhibitory) and EXP-1 (excitatory)) and metabotropic (GBB-1 and GBB-2) GABAB receptors in *C. elegans*. (**B** and **C**) Quantification (B) and DIC/fluorescent micrographs (C) of UPR^mt^ reporter (*hsp-6p::GFP*) expression in L4 larvae of wild-type, *gbb-1(tm1406), gbb-2(tm1165), gbb-1(tm1406*); *gbb-2(tm1165), unc-49(e407*) and *exp-1(ok1131*) animals. (**D**) Quantification of UPR^mt^ reporter (*hsp-6p::GFP*) expression in L4 larvae of wild-type, *gbb-1(tm1406), unc-25(e156*) and *unc-25(e156); gbb-1(tm1406*) animals. (**E**) Quantification of UPRmt reporter (*hsp-6p::GFP*) expression in L4 larvae of wild-type, *gbb-1(tm1406*) and *gbb-1(tm1406); xuEx1617[rgef-1p::gbb-1 cDNA]* animals. n = 30. *P* values assessed by one-way analysis of variance (ANOVA) Tukey’s post hoc test. Error bars indicate SEM. n.s., not significant. Scale bar, 250µm.

Within the *C. elegans* ventral nerve cord, D-type GABAergic motor neurons synapse with body wall muscle and adjacent cholinergic neurons, which are the only motor neurons that express *gbb-1* and *gbb-2* (Figure 3A) (*33, 34*). As GABA is generally an inhibitory neurotransmitter, and ACh is the primary excitatory neurotransmitter, these neurons work in a negative feedback loop to regulate each other and the muscles they innervate (*35*). When the inhibitory GABA signal is lost, cholinergic neurons are overactive, leading to increased ACh release (*34*). We therefore speculated that GABA regulates the intestinal UPR^mt^ through downstream ACh signalling. Our initial screening data showed that loss of the choline acetyltransferase CHA-1/ChAT, which is required for ACh production, significantly reduced intestinal *hsp-6p::gfp* expression (Figure 1A). Likewise, loss of the ACh vesicular transporter UNC-17/VAChT also reduced intestinal *hsp-6p::gfp* expression (Figure S4). These data support the concept that ACh levels can influence the systemic UPR^mt^, and that increased ACh release in animals lacking GABA signalling would increase the intestinal UPR^mt^. To test this hypothesis, we examined animals lacking two acetylcholinesterases, ACE-1/2, which have an approximately two-fold increase in systemic ACh (*36*). We found that, as with loss of GABA signalling, increased ACh in *ace-2(-); ace-1(-*) animals induced intestinal *hsp-6p::gfp* expression (Figure 3B-C).

**Fig. 3.**
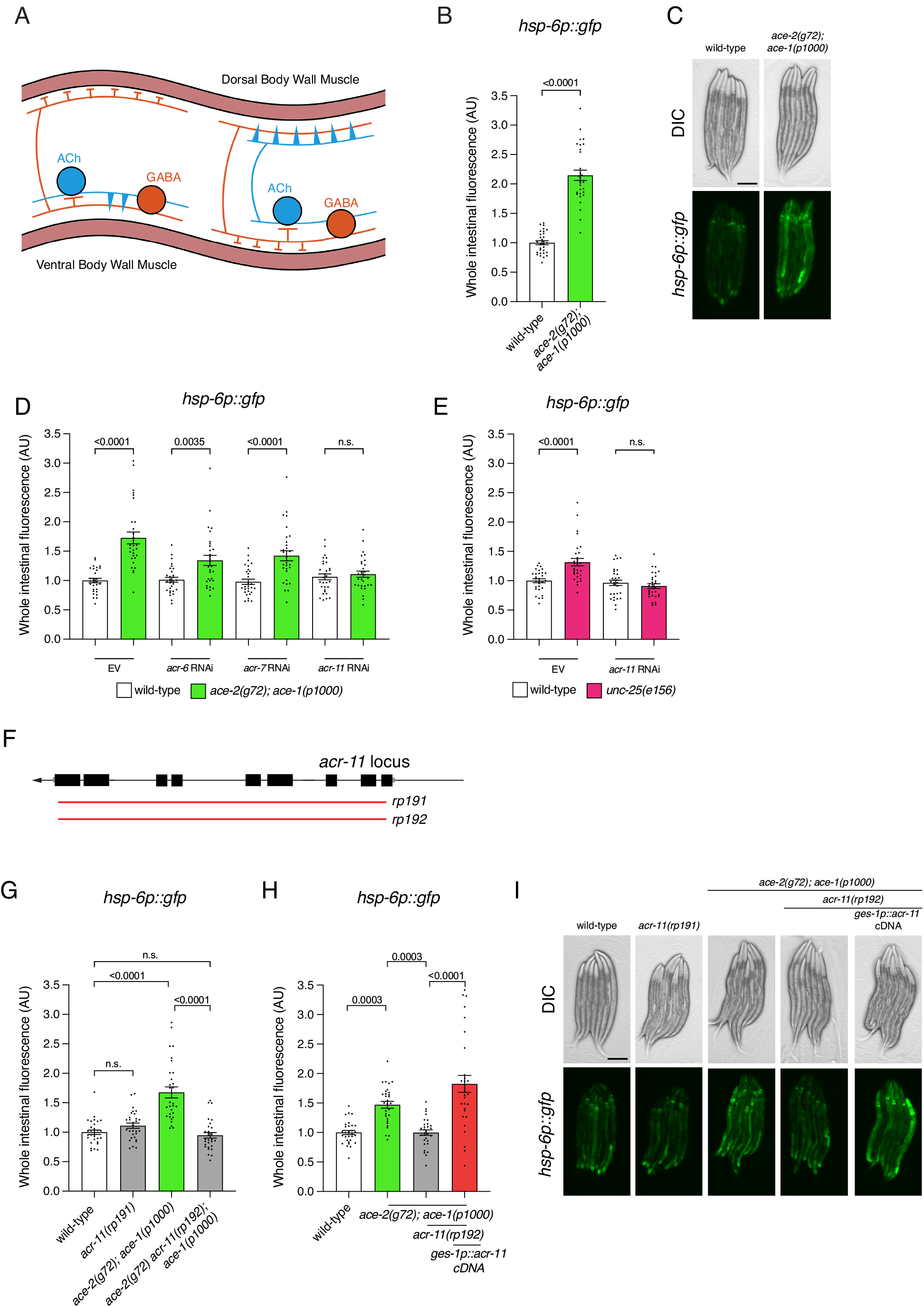
Acetylcholine promotes cell nonautonomous UPR^mt^ through the intestinal ACR-11/nicotinic α7 receptor. (**A**)Schematic of the *C. elegans* motor circuit. Orange = inhibitory GABAergic motor neurons; blue = excitatory cholinergic motor neurons; Pink = body wall muscle. (**B** and **C**) Quantification (B) and DIC/fluorescent micrographs (C) of UPR^mt^ reporter (*hsp-6p::GFP*) expression in L4 larvae of wild-type and *ace-2(g72); ace-1(p1000*) animals. (**D**) Quantification of UPR^mt^ reporter (*hsp-6p::GFP*) expression in L4 larvae of wild-type and *ace-2(g72); ace-1(p1000*) animals grown on empty vector (EV), *acr-6, acr-7* or *acr-11* RNAi from the mother’s L4 stage. (**E**)Quantification of UPR^mt^ reporter (*hsp-6p::GFP*) expression in L4 larvae of wild-type and *unc-25(e156*) animals grown on empty vector (EV) or *acr-11* RNAi from the mother’s L4 stage. (**F**)Structure of the *acr-11* locus. Black boxes, coding regions; grey boxes, untranslated regions; red lines, CRISPR/Cas9-generated deletion alleles. *rp191* = 3038bp deletion in wild-type animals; *rp192* = 3046bp deletion in *ace-2(g72); ace-1(p1000*) animals. (**G** to **I**) Quantification (G and H) and DIC/fluorescent micrographs (I) of UPR^mt^ reporter (*hsp-6p::GFP*) expression in L4 larvae of wild-type *acr-11(rp191), ace-2(g72); ace-1(p1000), ace-2(g72) acr-11(rp192); ace-1(p1000*) and *ace-2(g72) acr-11(rp192); ace-1(p1000); rpEx2314[ges-1p::acr-11 cDNA]* animals. n = 30. *P* values assessed by one-way analysis of variance (ANOVA) Tukey’s post hoc test (A and E) and unpaired t test with Welch’s correction (F and G). Error bars indicate SEM. n.s., not significant. Scale bars, 250µm.

We theorized that excess ACh acts directly on intestinal cells to induce the UPR^mt^. Therefore, we used RNAi to knock down intestinally-expressed ACh receptors in the *ace-2(-); ace-1(-*) compound mutant, screening for inhibition of *hsp-6p::gfp* induction (Figure 3D). We found that knockdown of the nicotinic α7 receptor (α7 nAChR) ACR-11, an ortholog of human CHRNA7, abrogated intestinal *hsp-6p::gfp* induction in *ace-2(-); ace-1(-*) animals (Figure 3D). ACR-11 knockdown also prevented *hsp-6p::gfp* induction in *unc-25(-*) animals (Figure 3E). We then used CRISPR/Cas9 to generate independent *acr-11* deletions in wild type and *ace-2(-); ace-1(-*) mutant animals (Figure 3F). As *acr-11* and *ace-2* are tightly linked on chromosome I, the *acr-11* deletions were generated separately. Confirming our RNAi data, the *acr-11(rp192*) mutation prevented *hsp-6p::gfp* induction in *ace-2(-); ace-1(-*) animals (Figure 3G,I). To examine the expression pattern of *acr-11*, we generated transgenic animals expressing green fluorescent protein (GFP) under control of the 2024bp *acr-11* promoter and detected expression in the intestine, and not in neurons or body wall muscle (Figure S5). To confirm the intestinal requirement of ACR-11, we resupplied *acr-11* cDNA in the intestine (*ges-1* promoter) to the *ace-2(-) acr-11(-); ace-1(-*) triple mutant, which restored the *hsp-6p::gfp* expression levels to that of the *ace-2(-); ace-1(-*) double mutant (Figure 3H-I). These data reveal that ACh acts through ACR-11 in the intestine to regulate the UPR^mt^.

Calcium plays key roles in mitochondrial function and health, including in stress response activation (*37, 38*). As ACR-11 is an α7 nAChR, which are highly permeable to calcium ions (*39*), we investigated intracellular calcium storage using an intestine-specific fluorescence resonance energy transfer (FRET)-based calcium indicator (*40, 41*) in animals with perturbed GABA and ACh signalling (Figure 4A-B). We found that both *unc-25(-*) and *ace-2(-); ace-1(-*) animals had increased intracellular calcium levels (Figure 4A-B). Furthermore, animals lacking ACR-11, either alone or when combined with the *ace-2(-); ace-1(-*) compound mutant, had wild-type calcium levels (Figure 4A-B). These data mirror *hsp-6p::gfp* reporter levels in these mutants, supporting our hypothesis that ACR-11 responds to ACh release by increasing intracellular calcium levels in the intestine, which in turn activates the UPR^mt^.

**Fig. 4.**
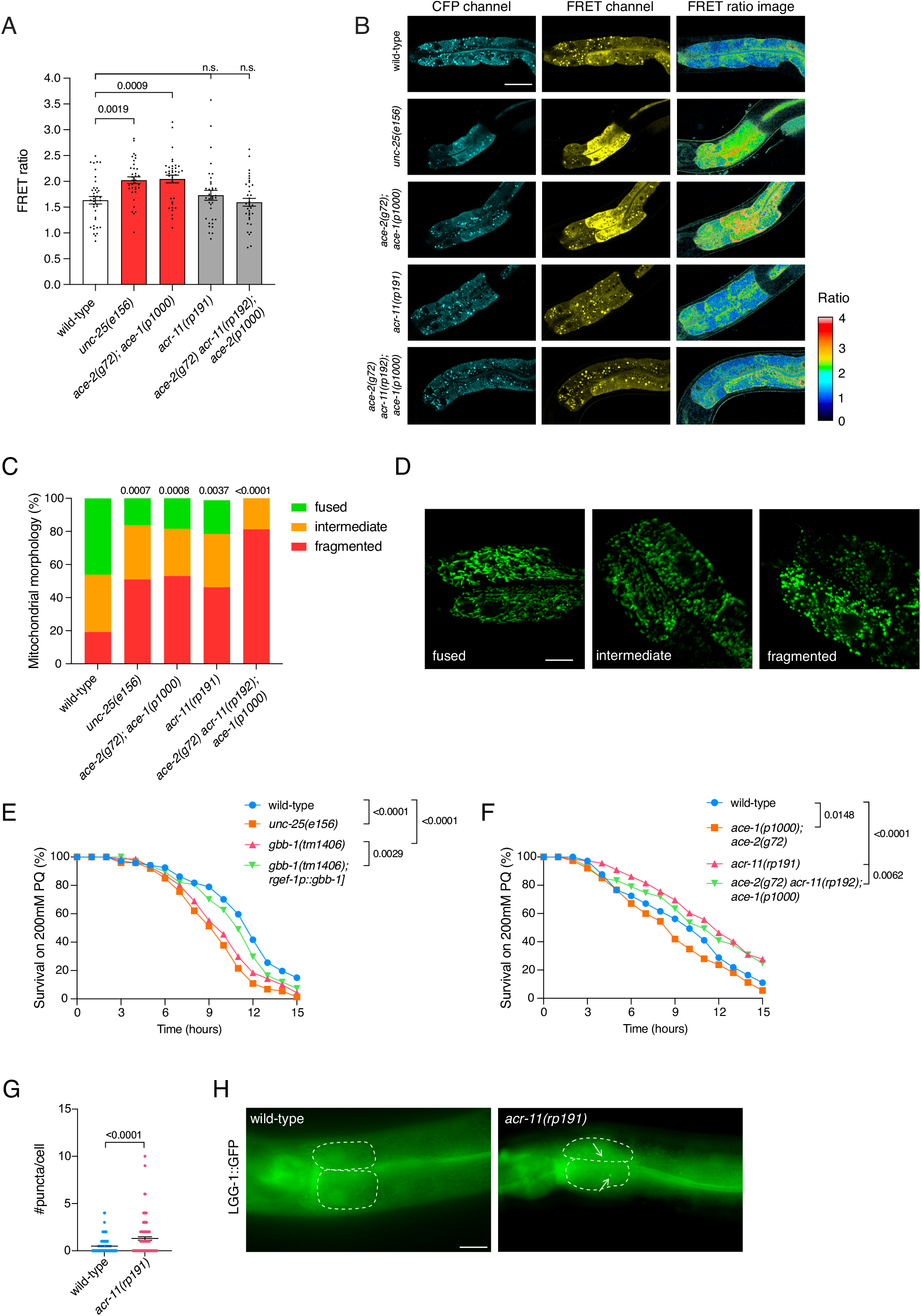
GABA/Ach regulation of ACR-11 influence mitochondrial dynamics and stress resistance. (**A** and **B**) Quantification (A) and micrographs (B) of calcium imaging by FRET microscopy of the two anterior intestinal cells of wild-type, *unc-25(e156), ace-2(g72); ace-1(p1000), acr-11(rp191*) and *ace-2(g72) acr-11(rp192); ace-1(p1000*) animals expressing *rnyEx109[nhx-2p::D3cpv]*, a ratiometric indicator. n = 38, 36, 36, 35, 36 (left to right). Scale bars, 50µm. (**C**) Survival analysis of wild-type, *unc-25(e156), gbb-1(tm1406*) and *gbb-1(tm1406); xuEx1617[rgef-1p::gbb-1 cDNA]* animals exposed to 200mM paraquat from the L4 larval stage. n = 67, 74, 70, 67 (top to bottom). (**D**) Survival analysis of wild-type, *unc-25(e156), ace-2(g72); ace-1(p1000), acr-11(rp191*) and *ace-2(g72) acr-11(rp192); ace-1(p1000*) animals exposed to 200mM paraquat from the L4 larval stage. n = 73, 72, 65, 67 (top to bottom). (**E** and **F**) Quantification (C) and confocal images (C) of mitochondrial morphology in the two anterior intestinal cells of wild-type, *unc-25(e156), ace-2(g72); ace-1(p1000), acr-11(rp191*) and *ace-2(g72) acr-11(rp192); ace-1(p1000*) animals expressing *zcIs17[ges-1p::GFP(mit)]*. Mitochondrial categorization: fused = tubular, intermediate = tubular and round, fragmented = round. N = 52, 49, 49, 54, 48 (left to right). Scale bars, 5µm. (**G** and **H**) Quantification (G) and micrographs (H) of GFP fluorescence in the two anterior intestinal cells of wild-type and *acr-11(rp191*) L4 larvae expressing GFP::LGG-1. n = 100. *P* values assessed by one-way analysis of variance (ANOVA) Tukey’s post hoc test (A), two-way analysis of variance (ANOVA) Tukey’s post hoc test (C and D), Chi square test (E) and unpaired t test (G). Error bars indicate SEM.

UPR^mt^ activation can have a positive or negative effect on mitochondrial health (*42*). An overactive stress response in the absence of stressors may prime mitochondria to manage subsequent stress exposure. Conversely, UPR^mt^ induction due to poor mitochondrial health may cause sensitivity to mitochondrial stressors. To assess whether perturbed GABA/ACh signalling influences mitochondrial stress resistance, we examined acute paraquat sensitivity. Paraquat exposure induces mitochondrial stress by disrupting complex I of the electron transport chain and increases superoxide levels (*43*). We found that loss of GABA biosynthesis (*unc-25* mutant) or metabotropic GABA receptors (*gbb-1* and *gbb-2* mutants) were sensitive to paraquat (Figure 4C and S6). This sensitivity was rescued in the *gbb-1(tm1406*) mutant by resupplying *gbb-1* cDNA in neurons (Figure 4C). Further, we found that *ace-2(-); ace-1(-*) animals were sensitive to paraquat exposure (Figure 4D). Thus, two conditions where ACh signalling is amplified causes sensitivity to a mitochondrial stressor. In contrast, *acr-11(-*) single and *ace-2(-) acr-11(-); ace-1(-*) triple mutant animals exhibited increased paraquat resistance (Figure 4D). As increased ACh levels (ACE-1/2 loss) is not additive to loss of ACR-11, this implies that loss of the ACR-11 α7 nAChR receptor provides mitochondrial stress resistance independent of ACh.

Another indicator of mitochondrial health is mitochondrial morphology. In general, more mitochondrial fusion indicates increased functionality, whereas more fragmentation indicates decreased functionality (*44*). Based on our paraquat data, we hypothesized that *unc-25(-*) and *ace-2(-); ace-1(-*) animals would have more fragmented mitochondria, and that *acr-11(-*) single and *ace-2(-) acr-11(-); ace-1(-*) triple mutants would have increased mitochondrial fusion. Using a mitochondrial-targeted GFP reporter expressed in the intestine (*ges-1p::gfp*^*mt*^) (*45*), we found that *unc-25(-*) and the *ace-2(-); ace-1(-*) animals had increased mitochondrial fragmentation in the intestine compared to wild type animals, as predicted (Figure 4E-F). This phenotype was intestine-specific, as we did not detect mitochondrial morphology changes in body wall muscle using a ubiquitously expressed reporter (*cox-4p::gfp*^*mt*^) in *unc-25(-*) animals (Figure S7) (*46*). Unexpectedly, the *acr-11(-*) single mutant also displayed a fragmented mitochondria phenotype (Figure 4E-F). Further, the *ace-2(-) acr-11(-); ace-1(-*) triple mutant exhibited more mitochondrial fragmentation than the *acr-11(-*) single and *ace-2(-); ace-1(-*) double mutants (Figure 4E-F). These data imply that, in the context of mitochondrial morphology, ACh signalling and the ACR-11 receptor act in distinct pathways.

We wondered how similar mitochondrial fragmentation phenotypes observed in *unc-25(-), ace-2(-); ace-1(-*) and *acr-11(-*) animals could lead to opposing mitochondrial fitness in terms of paraquat resistance. We posited that, in a pathway separate from GABA/ACh signalling, ACR-11 may repress mitophagy, meaning that ACR-11 loss would induce fragmentation as mitochondria are excessively cleared from the system. Using a GFP::LGG-1 reporter to measure autophagy (*47*), we indeed found that the number of intestinal autophagosomes in *acr-11(-*) L4 larvae is higher compared to control animals (Figure 4G-H). This suggests that animals lacking ACR-11 exhibit elevated mitochondrial turn-over in unstressed conditions that conveys a survival advantage when exposed to oxidative stress.

In summary, we discovered a neuro-intestinal circuit that regulates the UPR^mt^, mitochondrial dynamics, and organismal survival. By screening animals lacking specific neurotransmitters, we found that balanced GABAergic and cholinergic signalling is essential for cell-nonautonomous UPR^mt^ regulation. We found that GABA released from motor neurons controls the UPR^mt^ through two metabotropic GABAB receptor subunits that are expressed on, and inhibit, cholinergic motor neurons. Appropriate control of ACh levels is critical for UPR^mt^ regulation, as animals lacking the ACh-degradative enzymes exhibit a two-fold induction of the UPR^mt^. Further, we discovered that UPR^mt^ induction by ACh is dependent on ACR-11, an α7 nAChR receptor acting in the intestine. In addition to regulating the UPR^mt^, elevated ACh induces mitochondrial fragmentation and reduces survival of oxidative stress. Interestingly, we found that the ACR-11α7 nAChR receptor can act independently to ACh, with animals lacking ACR-11 exhibiting more intestinal mitochondrial fragmentation and mitophagy irrespective of ACh levels. These distinctions likely provide animals lacking ACR-11 their survival advantage in oxidative stress conditions. In vertebrates, nicotinic α7 receptors can act in the liver (some functions of which are performed by the *C. elegans* intestine) to promote cell survival (*48*), suggesting that mechanisms we describe here are phylogenetically conserved.

## Acknowledgments

We thank members of the Pocock laboratory for advice and comments on the manuscript. Imaging for this project was performed at Monash Microimaging. Some strains were provided by the *Caenorhabditis* Genetics Center (University of Minnesota), which is funded by the NIH Office of Research Infrastructure Programs (P40 OD010440) and National BioResource Project of Japan. We thank Shawn Xu and Jianfeng Liu for strains and plasmids.

## Author contributions

Conceptualization: RC, RP

Methodology: RC, WC, AH, BH, RP

Investigation: RC, WC, AH, BH, RP

Visualization: RC, WC, AH, BH

Funding acquisition: RP

Project administration: RP

Supervision: AH, RP

Writing – original draft: RC

Writing – review & editing: WC, AH, BH, RP

## Data and materials availability

All data is available in the main text or supplementary materials. In addition, Source Data are provided with this paper. There are no accession codes, unique identifiers, or weblinks in our study and no restrictions on data availability. Materials will be available upon request from the Pocock laboratory.

## Supplementary Materials

Materials and Methods

Figs. S1 to S7

Tables S1

References (49–51)

Data S1

