## Supplementary Information for "A neuro-intestinal circuit controls mitochondrial dynamics and stress resistance"

**The PDF file includes:**

Materials and Methods
Figs. S1 to S7
Table S1
References

**Other Supplementary Materials for this manuscript include the following:**

Data S1

### Materials and Methods

#### *C. elegans* culture

All *C. elegans* strains were grown at 20°C on NGM agar seeded with *Escherichia coli* OP50, unless otherwise stated. All mutant strains were backcrossed to N2 at least three times and maintained well fed for a minimum of two generations prior to analysis. Strains used in this study are listed in Table S1.

#### Fluorescence microscopy and quantification

Worms were anesthetised in 0.1ng/ml levamisole on a 5% agarose pad. Images for analysis were taken using a Zeiss AXIO Imager M2 upright fluorescence microscope and ZEN 2.0 software at 20X objective, unless otherwise specified. ImageJ software was used to determine whole intestinal fluorescence by isolating the intestine area and measuring integrated density (IntDensity). The corrected cellular fluorescence was calculated as: CTCF = Intestinal Integrated Density – (area * background MeanGrey)

#### RNA-mediated interference (RNAi) by feeding

HT115 bacteria containing control (L4440 plasmid) or experimental plasmids were cultured overnight in LB containing 50μg/ml ampicillin and seeded uniformly over plates containing IPTG (Isopropyl β- d-1-thiogalactopyranoside). Plates were dried at 37°C and brought to room temperature before worms were applied. For *atfs-1* and *ubl-5* RNAi experiments, five L4 hermaphrodites were picked to RNAi plates and the subsequent generation was scored at L4. For *dve-1* RNAi experiments, five L4 hermaphrodites were picked to NGM and allowed to grow for five days. Animals were then washed from the NGM plates using M9 and bleached for four minutes using a 1:1 ratio of bleach (White King Premium) and 5M NaOH to extract eggs. Extracted eggs were washed three times in M9 and filtered through a 40μm mesh before being applied to RNAi plates. Animals were grown to L4 and imaged as described above.

#### *C. elegans* expression constructs and transgenic strain generation

Reporter gene constructs were generated by PCR amplifying DNA elements and cloning into Fire vectors. Constructs were injected into young adult hermaphrodites using standard methods.

***acr-11p::gfp*** **reporter construct:**

A 2024bp sequence upstream of the *acr-11* start codon was amplified from *C. elegans* genomic DNA with forward (GAAATGAAATAAGCTTGCGAAGAGAGCGAGGAGG) and reverse (CCAATCCCGGGGATCCTTCAAAAAAATGTGGCTAAG) primers incorporating *HindIII-BamHI* restriction sites. The *HindIII-BamHI* digested promoter fragment was ligated into *HindIII-BamHI* digested pPD95.75 vector. The resultant *Pacr-11::gfp* plasmid was injected at 50 ng/µl, with 3 ng/µl of P*myo-2::mCherry* vector and 120 ng/µl bacterial DNA.

***ges-1p::acr-11*** **rescue construct:**

The 1386bp *acr-11* cDNA was amplified from a *C. elegans* cDNA library with forward (AGGACCCTTGGCTAGCATGATATTTAATCTAATTAATAG) and reverse (GATATCAATACCATGGTTAGGCAATAATATGAGG) primers incorporating *NheI-NcoI* restriction sites. The *Pges-1p-sphk-1* vector was digested with *NheI-NcoI* and the *acr-11* cDNA was inserted using the In-Fusion HD Cloning Kit (Takara Bio) to replace the *sphk-1* sequence. The resultant *Pges-1::acr-11cDNA* plasmid was injected at 10 ng/µl, with 50 ng/µl of *Pttx-3::mCherry* vector and 120 ng/µl bacterial DNA.

***unc-25p::unc-25*** **rescue construct:**

A 1892bp sequence upstream of the *unc-25* start codon was amplified from *C. elegans* genomic DNA with forward (CGACTCTAGAGGATCCTGAGAAATAAGAAATAATTG) and reverse (CCAATCCCGGGGATCCTTTTTGGCGGTGAACTGAGC) primers incorporating *BamHI* restriction sites. The *BamHI* digested promoter fragment was ligated into the *BamHI* digested pPD49.26 vector.

The 1718bp *unc-25* sequence was amplified from *C. elegans* genomic DNA with forward (TGGCTAGCGTCGACGGTACCCCAAAAATGTCCTCTGCTG) and reverse (GATATCAATACCATGGTACCTATAAAAAAGTGTCGTATTCACTAC) primers incorporating *NheI* restriction sites. The *NheI* digested *unc-25* fragment was ligated into the *NheI* digested pPD49.26 + *unc-25p* vector.

The resultant *Punc-25p::unc-25* plasmid was injected at 5ng/µl, with 3ng/µl, of P*myo-2::mCherry* vector and 120 ng/µl bacterial DNA.

**CRISPR-Cas9 genome editing**

To generate *acr-11* deletion mutants, adult wild-type or *ace-2(p1000)I; ace-1(g72)X* hermaphrodites were microinjected with Cas9 protein and two crRNAs: 5' crRNA (CCTGTGCGACGGAAGTGTTG), 3' crRNA (CGAATCTCCAATCCGTTTGA). *acr-11* deletions were identified by PCR and confirmed by Sanger sequencing. The *acr-11(rp191)* allele generated in wild-type animals is a 3038bp deletion and the *acr-11(rp192)* allele generated in *ace-2(g72)I; ace-1(p1000)X* animals is a 3046bp deletion. Both deletion alleles remove most of the *acr-11* gene (Figure xx). Each allele was backcrossed to wild-type males prior to analysis.

**Calcium imaging**

Calcium levels were visualised using the calcium indicator d3cpv expressed in the intestine using the *nhx-2* intestine-limited promoter as previously described (*1*). Worms were anesthetised with in 0.1ng/ml levamisole diluted in M9 and mounted to 5% agarose pads. Images were taken using Leica Stellaris5 Invert Confocal Microscope and objective lens 63x/1.40 Oil (WD 140 μm). CFP (405 excitation, 455-495 emission) and FRET (405 excitation, 515-555 emission) were collected. The FRET ratio in the first three intestinal cell pairs was calculated as (FRETint – FRETbg) / (CFPint – CFPbg).

**Mitochondrial morphology analysis**

Worms were anesthetised with 50mM sodium azide diluted in M9 and mounted to 5% agarose pads. Images were taken using (Leica Stellaris5 Invert Confocal Microscope and objective lens 63x/1.40 Oil (WD 140 μm). Optical slice thickness was 0.2μm. Z stack images were blinded to genotype and mitochondrial morphology classified as fused, intermediate or fragmented (*2*).

**Acute paraquat sensitivity assays**

200mM paraquat plates were prepared as described previously (*3*) and seeded with 50μl 10x concentrated OP50. 25 L4 worms were transferred to each plate and survival was assessed each hour for 15 consecutive hours. Worms that left the agar were excluded from analysis.

### Mitophagy analysis

Intestinal mitophagy was analysed using the *lgg-1p::gfp::lgg-1* reporter. A single image was taken in line with the nucleus of the first two intestinal cells using a Zeiss AXIO Imager M2 upright fluorescence microscope as described above. Images were blinded and LGG-1 puncta were counted.

### Statistical analysis

Statistical analysis was performed using GraphPad Prism 9 software. Statistical tests and n numbers are indicated in corresponding figure legends. Differences with a p-value <0.05 were considered significant.

**Supplementary Figures**

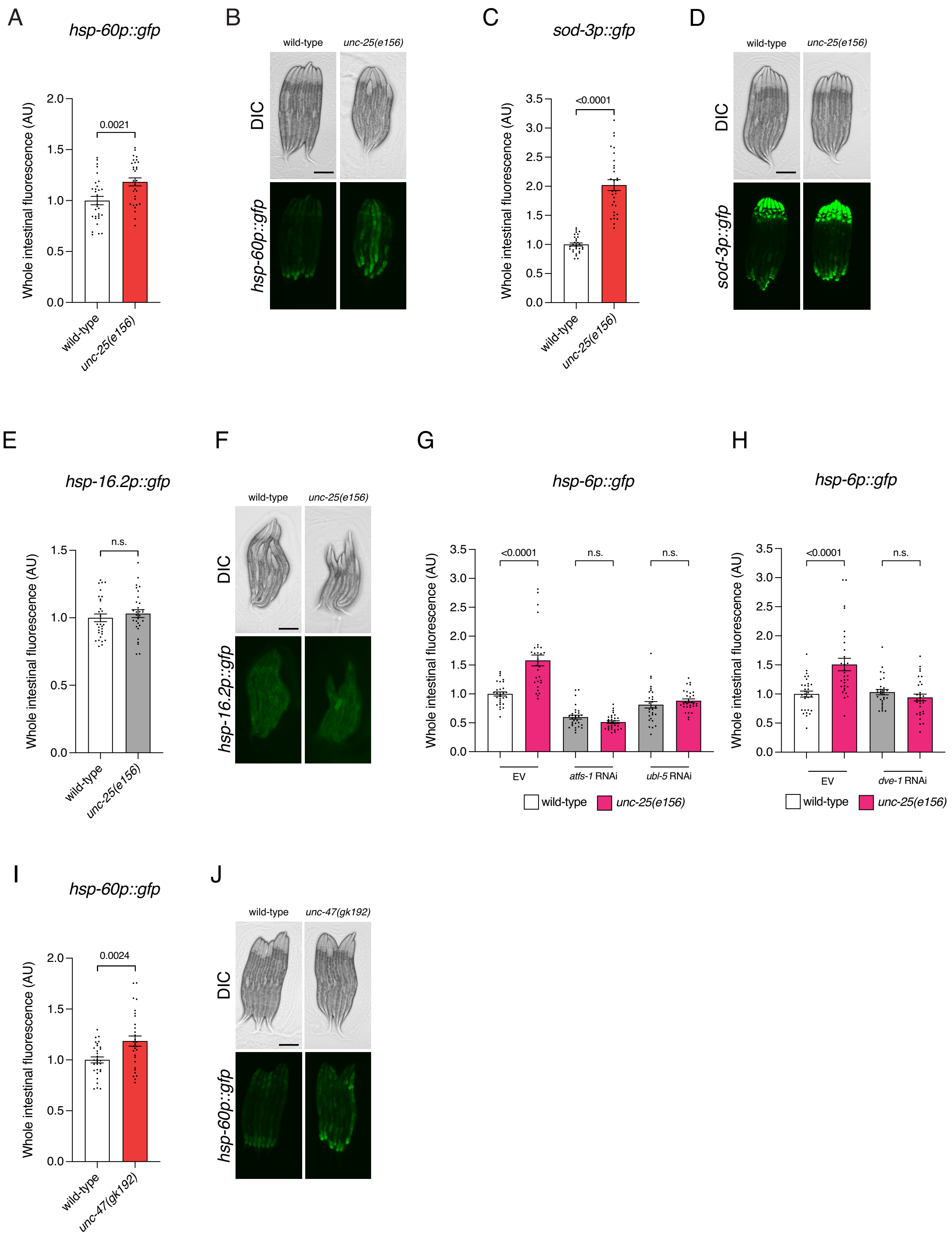

A
hsp-60p::gfp
Whole intestinal fluorescence (AU)
0.0021
wild-type
unc-25(e156)
B
wild-type
unc-25(e156)
DIC
hsp-60p::gfp
C
sod-3p::gfp
Whole intestinal fluorescence (AU)
<0.0001
wild-type
unc-25(e156)
D
wild-type
unc-25(e156)
DIC
sod-3p::gfp
E
hsp-16.2p::gfp
Whole intestinal fluorescence (AU)
n.s.
wild-type
unc-25(e156)
F
wild-type
unc-25(e156)
DIC
hsp-16.2p::gfp
G
hsp-6p::gfp
Whole intestinal fluorescence (AU)
<0.0001
n.s.
n.s.
EV
atfs-1 RNAi
ubl-5 RNAi
wild-type
unc-25(e156)
H
hsp-6p::gfp
Whole intestinal fluorescence (AU)
<0.0001
n.s.
EV
dve-1 RNAi
wild-type
unc-25(e156)
I
hsp-60p::gfp
Whole intestinal fluorescence (AU)
0.0024
wild-type
unc-47(gk192)
J
wild-type
unc-47(gk192)
DIC
hsp-60p::gfp

**Fig. S1. GABA regulation of stress signalling.**
**(A** and **B)** Quantification (A) and DIC/fluorescent micrographs (B) of $UPR^{mt}$ reporter (*hsp-60p::GFP*) expression in L4 larvae of wild-type and *unc-25(e156)* animals.
**(C** and **D)** Quantification (C) and DIC/fluorescent micrographs (D) of oxidative stress reporter (*sod-3p::GFP*) expression in L4 larvae of wild-type and *unc-25(e156)* animals.
**(E** and **F)** Quantification (E) and DIC/fluorescent micrographs (F) of cytosolic heat stress reporter (*hsp-16.2p::GFP*) expression in L4 larvae of wild-type and *unc-25(e156)* animals.
**(G** and **H)** Quantification of $UPR^{mt}$ reporter (*hsp-6p::GFP*) expression in L4 larvae of wild-type and *unc-25(e156)* animals grown on empty vector (EV), *atfs-1*, *ubl-5* or *dve-1* RNAi from the mother's L4 stage.
(**I**) Quantification (I) and DIC/fluorescent micrographs (J) of $UPR^{mt}$ (*hsp-60p::GFP*) expression in L4 larvae of wild-type and *unc-47(gk192)* animals.
n = 30. *P* values assessed by unpaired t test with Welch's correction (A, C, E and I) or one-way analysis of variance (ANOVA) Tukey's post hoc test (G and H). Error bars indicate SEM. n.s., not significant. Scale bars, 250µm.

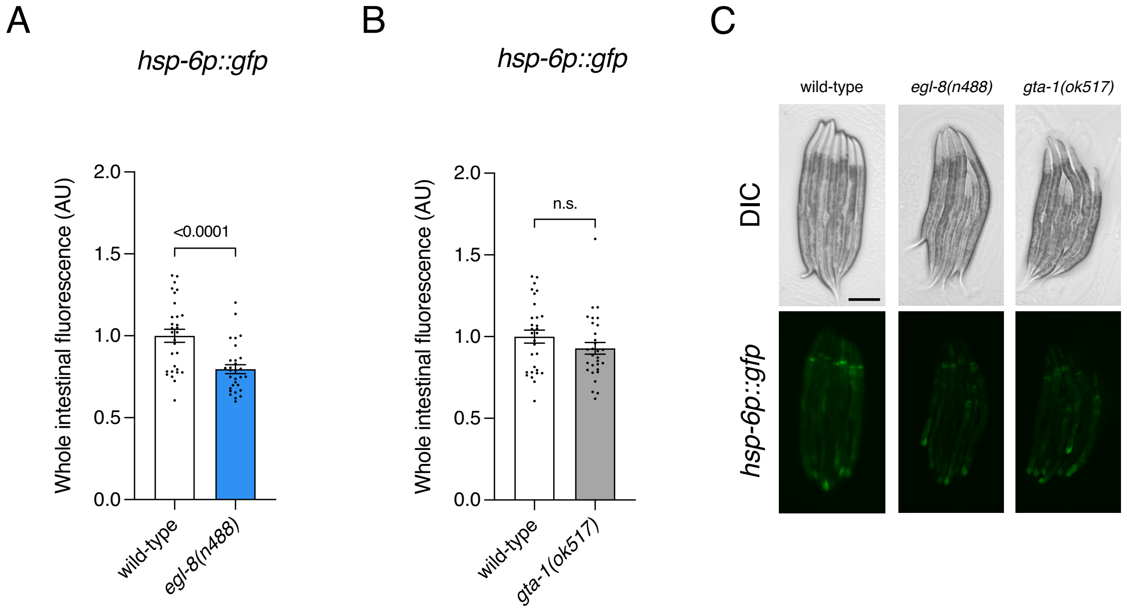


**Fig. S2. GABA regulates the UPR$^{mt}$ independently of the GABA shunt and longevity pathways.** (**A** to **C**) Quantification (A-B) and DIC/fluorescent micrographs (C) of UPR$^{mt}$ reporter (*hsp-6p::GFP*) expression in L4 larvae of wild-type, *egl-8(n488)* (A) and *gta-1(ok517)* (B) animals.
n = 30. *P* values assessed by unpaired t test with Welch's correction. Error bars indicate SEM. n.s., not significant. Scale bar, 250µm.

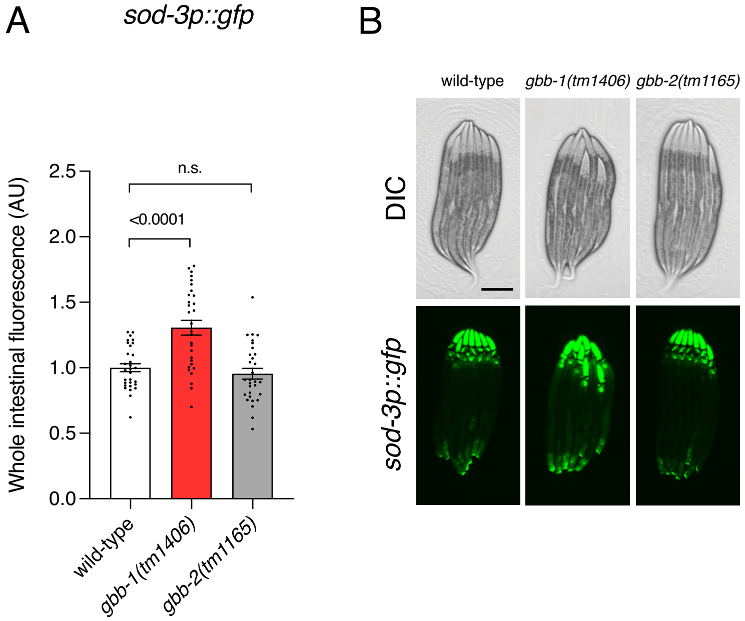


**Fig. S3. An oxidative stress reporter is regulated by GBB-1 and not GBB-2.**
**(A and B)** Quantification (A) and DIC/fluorescent micrographs (B) of oxidative stress reporter (*sod-3p::GFP*) expression in L4 larvae of wild-type, *gbb-1(tm1406)* and *gbb-2(tm1165)* animals.
n = 30. *P* values assessed by one-way analysis of variance (ANOVA) Tukey's post hoc test. Error bars indicate SEM. n.s., not significant. Scale bar, 250µm.

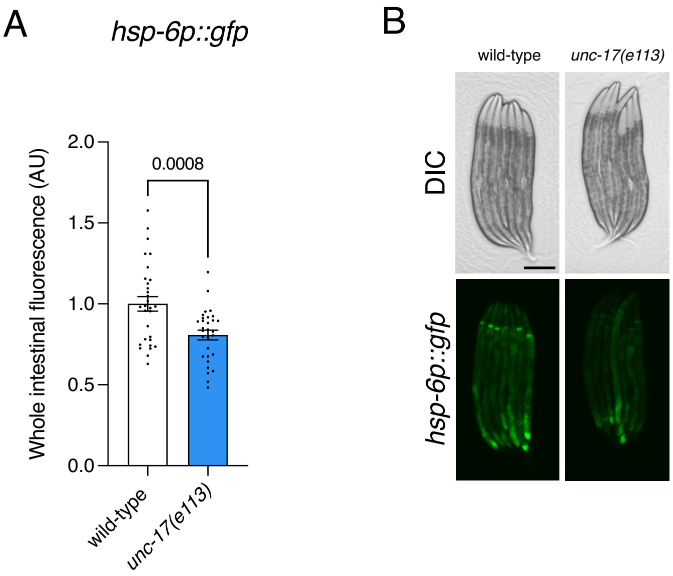


**Fig. S4. The ACh vesicular transporter UNC-17/VAChT is required for UPR$^{mt}$ reporter expression.**
**(A** and **B)** Quantification (A) and DIC/fluorescent micrographs (B) of UPR$^{mt}$ reporter (*hsp-6p::GFP*) expression in L4 larvae of wild-type and *unc-17(e113)* animals.
n = 30. *P* values assessed by unpaired t test with Welch's correction. Error bars indicate SEM. n.s., not significant. Scale bar, 250μm.

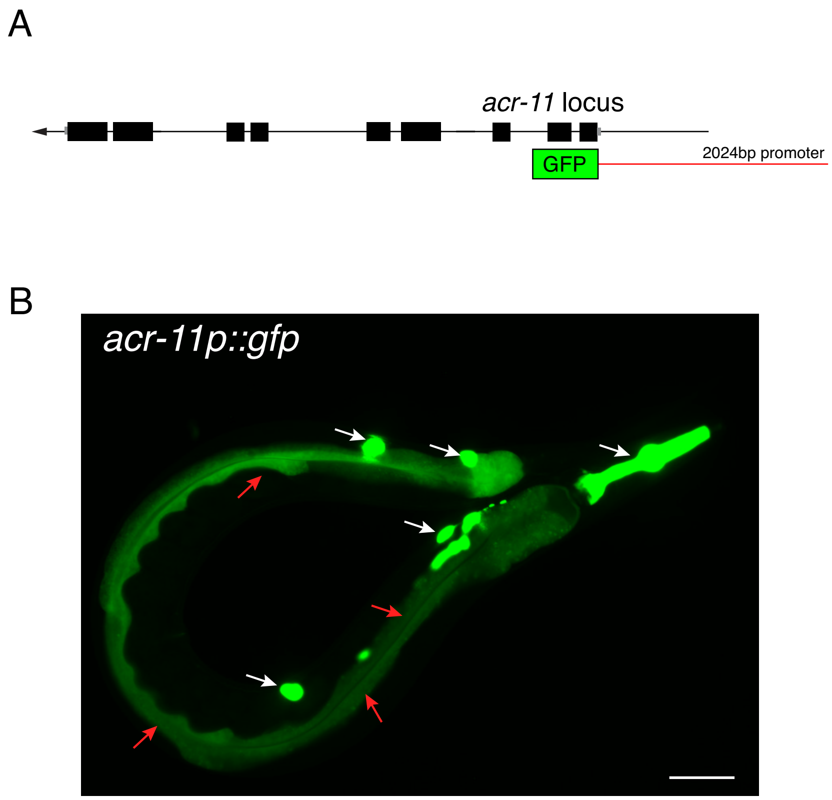


**Fig. S5. The *acr-11* promoter drives expression in the intestine.**
(**A** to **B**) Structure of the *acr-11* locus. Black boxes, coding regions; grey boxes, untranslated regions; red line, 2024bp *acr-11* upstream sequence used to generate the *acr-11p::GFP* transcriptional reporter (A). Fluorescent micrograph (B) of a transgenic animal expressing *acr-11p(2024bp)::GFP*. Red arrows = intestinal expression; white arrows = background pharyngeal and coelomocyte expression from the *myo-2::mCherry* co-injection marker. Scale bar, 25µm.

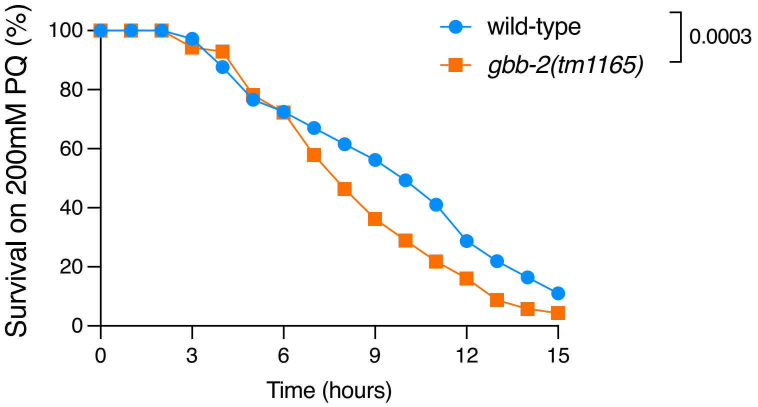


**Fig. S6. The GBB-2 GABA$_B$ receptor is required for resistance to oxidative stress.**
Survival analysis of wild-type and *gbb-2(tm1165)* animals exposed to 200mM paraquat from the L4 larval stage. n = 69, 73 (left to right). *P* values assessed by two-way analysis of variance (ANOVA) Tukey's post hoc test.

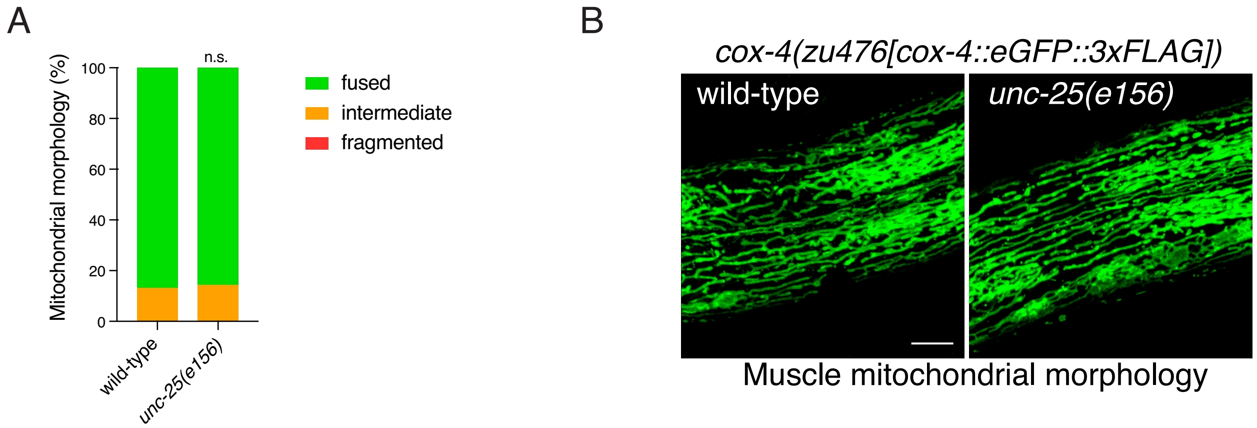


**Fig. S7. GABA does not control mitochondrial morphology in body wall muscle.**
(**A** and **B**) Quantification (A) and confocal images (B) of mitochondrial networks in body wall muscle cells of wild-type and *unc-25(e156)* animals expressing *cox-4(zu476)[cox-4::eGFP::3xFLAG])*. n = 21, 23 (left to right). *P* value assessed by Chi-square test.

**Table S1.** ***C. elegans*** **strains used in this work.**

| Strain name | Genotype | Source |
|---|---|---|
| Bristol strain N2 | wild type | CGC |
| SJ4100 | *zcIs13[hsp-6p::GFP + lin-15(+)]V* | CGC |
| RJP5029 | *cat-2(n4547)II; zcIs13[hsp-6p::GFP + lin-15(+)]V* | this study |
| RJP5000 | *tdc-1(n3420)II; zcIs13[hsp-6p::GFP + lin-15(+)]V* | this study |
| RJP4817 | *tph-1(mg280)II; zcIs13[hsp-6p::GFP + lin-15(+)]V* | this study |
| RJP4815 | *eat-4(ky5)III; zcIs13[hsp-6p::GFP + lin-15(+)]V* | this study |
| RJP4821 | *unc-25(e156)III; zcIs13[hsp-6p::GFP + lin-15(+)]V* | this study |
| RJP5021 | *cha-1(p1152)IV; zcIs13[hsp-6p::GFP + lin-15(+)]V* | this study |
| RJP5762 | *unc-25(e156)III; zcIs13[hsp-6p::GFP + lin-15(+)]V; rpEx2310[unc-25p::unc-25 cDNA]* | this study |
| RJP5242 | *unc-47(gk192)III; zcIs13[hsp-6p::GFP + lin-15(+)]V* | this study |
| RJP5110 | *unc-30(e191)IV; zcIs13[hsp-6p::GFP + lin-15(+)]V* | this study |
| SJ4058 | *zcIs9[hsp-60p::GFP + lin-5(+)]V* | CGC |
| RJP5002 | *unc-25(e156)III; zcIs9[hsp-60p::GFP + lin-5(+)]V* | this study |
| CF1553 | *muIs84[sod-3p::GFP + rol-6(su1006)]* | CGC |
| RJP5291 | *unc-25(e156)III; muIs84[sod-3p::GFP + rol-6(su1006)]* | this study |
| CL2070 | *dvIs70[hsp-16.2p::GFP + rol-6(su1006)]* | CGC |
| RJP5651 | *unc-25(e156)III; dvIs70[hsp-16.2p::GFP + rol-6(su1006)]* | this study |
| RJP5307 | *unc-47(gk192)III; zcIs9[hsp-60p::GFP + lin-5(+)]V* | this study |
| CB156 | *unc-25(e156)III* | CGC |
| RJP5240 | *zcIs13[hsp-6p::GFP + lin-15(+)]V; gbb-1(tm1406)X* | this study |
| RJP5241 | *gbb-2(tm1165)IV; zcIs13[hsp-6p::GFP + lin-15(+)]V* | this study |
| RJP5449 | *gbb-2(tm1165)IV; zcIs13[hsp-6p::GFP + lin-15(+)]; gbb-1(tm1406)X* | this study |
| RJP5652 | *gbb-1(tm1406)X; muIs84[sod-3p::GFP + rol-6(su1006)]* | this study |
| RJP5653 | *gbb-2(tm1165)IV; muIs84[sod-3p::GFP + rol-6(su1006)]* | this study |
| RJP5564 | *zcIs13[hsp-6p::GFP + lin-15(+)]V; gbb-1(tm1406)X; xuEx1617[rgef-1p::gbb-1::SL2::mCherry]* | this study |
| | *zcIs13[hsp-6p::GFP + lin-15(+)]V; gbb-1(tm1406)X* | this study |
| RJP5447 | *gta-1(ok517)IV; zcIs13[hsp-6p::GFP + lin-15(+)]V* | this study |
| RJP5448 | *egl-8(n488)V; zcIs13[hsp-6p::GFP + lin-15(+)]V* | this study |
| RJP4884 | *unc-49(e407)III; zcIs13[hsp-6p::GFP + lin-15(+)]V* | this study |
| RJP5648 | *exp-1(ok1131)II; zcIs13[hsp-6p::GFP + lin-15(+)]V* | this study |
| RJP5589 | *unc-25(e156)III; zcIs13[hsp-6p::GFP + lin-15(+)]V; gbb-1(tm1406)X* | this study |
| RJP5833 | *unc-17(e113)IV; zcIs13[hsp-6p::GFP + lin-15(+)]V* | this study |
| RJP5743 | *ace-2(g72)I; zcIs13[hsp-6p::GFP + lin-15(+)]V; ace-1(p1000)X* | this study |

| | | |
|---|---|---|
| RJP5850 | *acr-11(rp191)I; zcIs13[hsp-6p::GFP + lin-15(+)]V* | this study |
| RJP5851 | *acr-11(rp192)I; ace-2(g72)I; zcIs13[hsp-6p::GFP + lin-15(+)]V; ace-1(p1000)X* | this study |
| RJP5875 | *acr-11(rp192)I; ace-2(g72)I; zcIs13[hsp-6p::GFP + lin-15(+)]V; ace-1(p1000)X rpEx2314[ges-1p::acr-11 cDNA]* | this study |
| RJP5832 | *ace-2(g72)I; gbb-2(tm1165)IV; ace-1(p1000)X; zcIs13[hsp-6p::GFP + lin-15(+)]V* | this study |
| RJP5902 | *rpEx2313[acr-11p::GFP]* | this study |
| TQ37C | *gbb-1(tm1406)X* | JianFeng Liu |
| TQ5123 | *gbb-1(tm1406)X; xuEx1617[rgef-1p::gbb-1::SL2::mCherry]* | JianFeng Liu |
| GG201 | *ace-2(g72)I; ace-1(p1000)X* | CGC |
| RJP5825 | *acr-11(rp191)I* | this study |
| RJP5826 | *acr-11(rp192)I; ace-2(g72)I; ace-1(p1000)X* | this study |
| SJ4143 | *zcIs17[ges-1p::GFP(mit)]* | CGC |
| RJP5318 | *unc-25(e156)III; zcIs17[ges-1p::GFP(mit)]* | this study |
| RJP5831 | *ace-2(g72)I; ace-1(p1000)X; zcIs17[ges-1p::GFP(mit)]* | this study |
| RJP5852 | *acr-11(rp191)I; zcIs17[ges-1p::GFP(mit)]* | this study |
| RJP5853 | *acr-11(rp192)I; ace-2(g72)I; ace-1(p1000)X; zcIs17[ges-1p::GFP(mit)]* | this study |
| JJ2586 | *cox-4(zu476[cox-4::eGFP::3xFLAG])I* | CGC |
| RJP5440 | *cox-4(zu476[cox-4::eGFP::3xFLAG])I; unc-25(e156)III* | this study |
| RJP5854 | *rnyEx109[nhx-2p::D3cpv + pha-1(+)]* | this study – derived from KWN190 |
| RJP5855 | *unc-25(e156)III; rnyEx109[nhx-2p::D3cpv + pha-1(+)]* | this study |
| RJP5856 | *ace-2(g72)I; ace-1(p1000)X; rnyEx109[nhx-2p::D3cpv + pha-1(+)]* | this study |
| RJP5857 | *acr-11(rp191)I; rnyEx109[nhx-2p::D3cpv + pha-1(+)]* | this study |
| RJP5858 | *acr-11(rp192)I; ace-2(g72)I; ace-1(p1000)X; rnyEx109[nhx-2p::D3cpv + pha-1(+)]* | this study |
| RJP5859 | *zcIs13[hsp-6p::GFP + lin-15(+)]V; rmIs110[rgef-1p::Q40::YFP]* | this study |
| RJP5860 | *acr-11(rp191)I; zcIs13[hsp-6p::GFP + lin-15(+)]V; rmIs110[rgef-1p::Q40::YFP]* | this study |
| TQ4911 | *gbb-2(tm1165)IV* | JianFeng Liu |
| RJP5874 | *gbb-2(tm1165)IV; gbb-1(tm1406)X* | this study |
| MAH236 | *sqIs13 [lgg-1p::GFP::lgg-1 + odr-1p::RFP]* | CGC |
| RJP5884 | *acr-11(rp191)I; sqIs13 [lgg-1p::GFP::lgg-1 + odr-1p::RFP]* | this study |

**Data S1. (separate file)**
Source Data.
